# SIGAR: Inferring features of genome architecture and DNA rearrangements by split read mapping

**DOI:** 10.1101/2020.05.05.079426

**Authors:** Yi Feng, Leslie Y. Beh, Wei-Jen Chang, Laura F. Landweber

## Abstract

Ciliates are microbial eukaryotes with distinct somatic and germline genomes. Post-zygotic development involves extensive remodeling of the germline genome to form somatic chromosomes. Ciliates therefore offer a valuable model for studying the architecture and evolution of programmed genome rearrangements. Current studies usually focus on a few model species, where rearrangement features are annotated by aligning reference germline and somatic genomes. While many high-quality somatic genomes have been assembled, a high quality germline genome assembly is difficult to obtain due to its smaller DNA content and abundance of repetitive sequences. To overcome these hurdles, we propose a new pipeline SIGAR (**<u>S</u>**plitread **I**nference of **<u>G</u>**enome **<u>A</u>**rchitecture and **<u>R</u>**earrangements) to infer germline genome architecture and rearrangement features without a germline genome assembly, requiring only short germline DNA sequencing reads. As a proof of principle, 93% of rearrangement junctions identified by SIGAR in the ciliate *Oxytricha trifallax* were validated by the existing germline assembly. We then applied SIGAR to six diverse ciliate species without germline genome assemblies, including *Ichthyophthirius multifilii,* a fish pathogen. Despite the high level of somatic DNA contamination in each sample, SIGAR successfully inferred rearrangement junctions, short eliminated sequences and potential scrambled genes in each species. This pipeline enables pilot surveys or exploration of DNA rearrangements in species with limited DNA material access, thereby providing new insights into the evolution of chromosome rearrangements.

## Introduction

Ciliates are model organisms for studying genome rearrangement. They exhibit nuclear dimorphism: each cell contains a somatic macronucleus (MAC) and a germline micronucleus (MIC). The MAC consists of high-copy number chromosomes that are transcriptionally active in vegetative growth. In contrast, the MIC genome is inert, and only involved in sexual conjugation. After mating, a new MAC genome rearranges from a copy of the zygotic MIC, together with massive DNA elimination (Chen et al. 2014; Hamilton et al. 2016). The retained, macronuclear-destined sequences (MDS) must be properly ordered and oriented, and sometimes even descrambled (Chen et al. 2014; Sheng et al. 2020), to form functional MAC chromosomes (Figure 1A).

**Figure 1.**
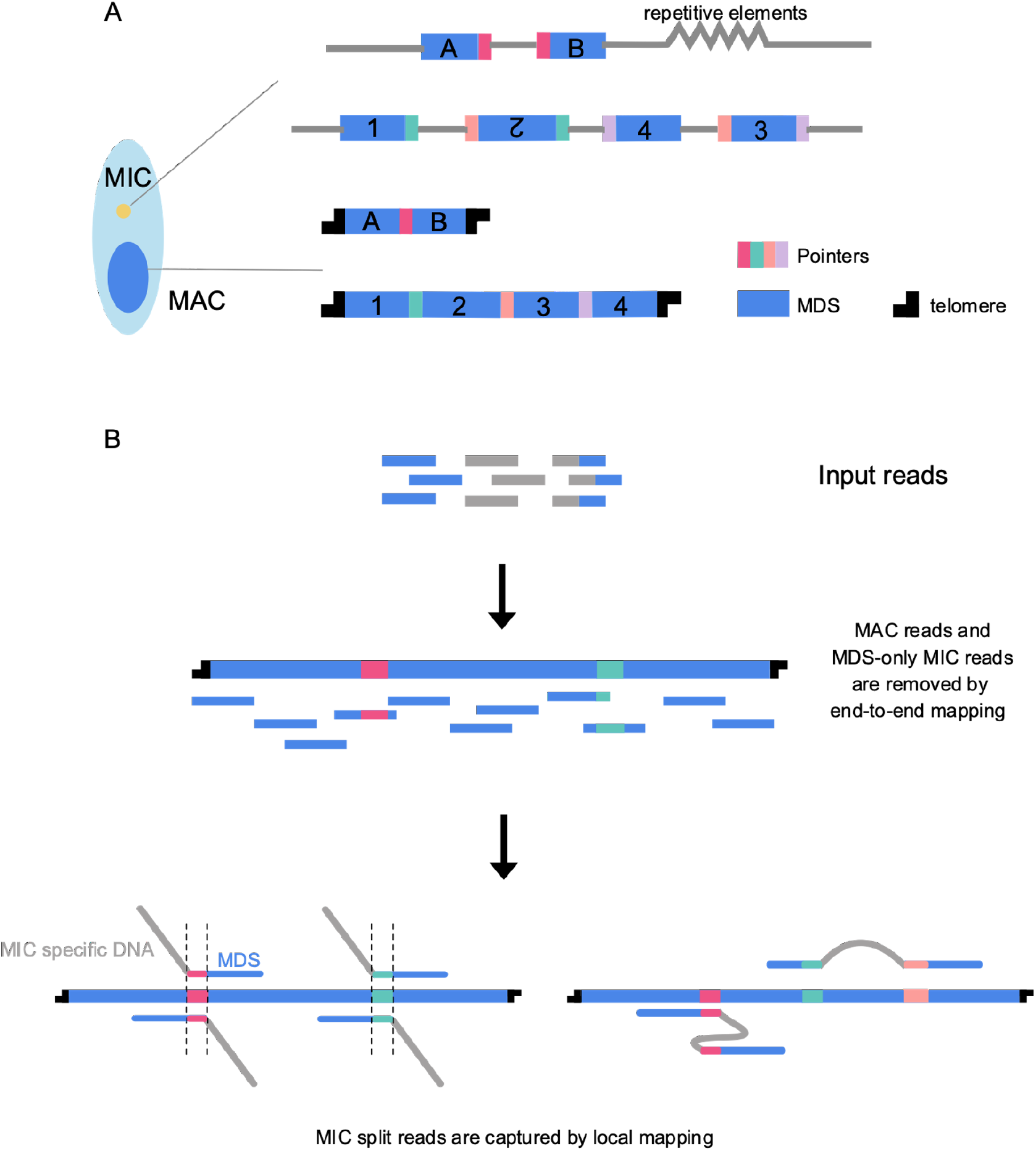
Schematic of genome rearrangements in ciliates and SIGAR strategy. **A)** Ciliates have separate germline (MIC) and somatic (MAC) genomes. The MAC chromosomes form from MIC DNA during development by elimination of intervening DNA sequences (gray) and reorganization of MDSs (blue). Some rearrangements join neighboring MDSs (e.g. A and B), whereas scrambled rearrangements require translocation and/or inversion (e.g. 1-4). Microhomologous pointer sequences, shown in different colors, are present at the end of MDS n and the beginning of MDS n+1 on the MIC chromosome, with one copy retained in the MDS-MDS junction on the MAC chromosome. **B)** SIGAR strategy. Reads that only contain MDS are removed by end-to-end mapping to MAC contigs. The filtered reads are mapped locally to identify MIC reads that split at MDS-MDS junctions. Such reads that map to both MDS n and MDS n+1 permit inference of the pointer sequence from the overlapped region and the eliminated sequence between them. Scrambled features of the germline map can sometimes be inferred from reads containing at least two mapped blocks.

Most genome rearrangement studies focus on model organisms like *Tetrahymena* (Hamilton et al. 2016), *Paramecium* (Guérin et al. 2017) and *Oxytricha* (Chen et al. 2014), which possess well-assembled MIC and MAC reference genomes for annotation of DNA rearrangements. Recent years have seen a bloom of *de novo* MAC genome assemblies in diverse ciliate species, including *Stentor* (Slabodnick et al. 2017), *Euplotes* (Wang et al. 2016; Chen et al. 2019), hypotrich ciliates (Chen et al. 2015) and *Tetrahymena* genus species (Xiong et al. 2019). MAC chromosomes are generally significantly shorter than MIC chromosomes, with some species exhibiting gene-sized “nanochromosomes”. Thus, many high quality MAC genomes have been assembled using short next-generation sequencing reads. By taking advantage of third generation sequencing long reads, some MAC genomes have been assembled with unprecedented completeness (Wang et al. 2020; Sheng et al. 2020), sometimes obviating the need for assembly when the average read length exceeds the typical chromosome length (Lindblad et al. 2019). In contrast, sequencing and assembling the long MIC genomes is experimentally and computationally complex. Besides having Mbp scale chromosomes at much lower copy number, the MIC also contains repetitive elements and centromeric regions that can best be resolved by third generation long reads. Purification of MIC genomic DNA that is free of MAC contamination is also a challenge. The MAC to MIC DNA ratio in cells ranges from 46 to 800 (Prescott 1994), which means that only 0.1-2% of the DNA in whole cells originates from the MIC. There do exist experimental methods to separate MIC and MAC nuclei, e.g. sucrose gradient centrifugation (Chen et al. 2014) and flow cytometry (Guérin et al. 2017). However, these techniques were developed for specific model ciliates, and are not generalizable across diverse species. Moreover, some ciliates are not free-living in nature (Coyne et al. 2011), or uncultivatable in the lab. The difficulty of obtaining high quality MIC-enriched DNA from these species presents additional obstacles to understanding germline genome architecture. Single-cell techniques can be helpful to analyze germline scaffolds, but require whole genome and transcriptome assemblies (Maurer-Alcalá et al. 2018).

To overcome these challenges and provide insight into germline genome architecture in the absence of a fully-assembled MIC genome, we propose a new pipeline, SIGAR (**<u>S</u>**plit-read **I**nference of **<u>G</u>**enome **<u>A</u>**rchitecture and **<u>R</u>**earrangements) using economical short, next generation reads and MAC genome assemblies. Rather than using a MIC assembly, SIGAR takes advantage of short MIC reads whose alignment diverges at MDS-MDS junctions in MAC chromosomes (Figure 1B). Here we validate SIGAR by showing high concordance of its results with published *Oxytricha* MIC genome annotations (Burns et al. 2016). We then used SIGAR to infer rearrangement features in 5 hypotrich ciliates and *Ichthyophthirius multifiliis,* yielding novel insights into MIC genome architecture in diverse phylogenetic lineages, all without genome assembly. This new pipeline will promote the use of published datasets to reveal more cases of DNA rearrangement, and offers the possibility to explore germline genome evolution in diverged ciliate species and natural isolates.

## Results

### SIGAR strategy

SIGAR infers MIC genome structure and DNA rearrangement features by identifying short MIC reads that partially map to MAC chromosomes. It first removes MDS-only reads by end-to-end mapping to enrich for MIC-specific reads in the dataset (Figure 1B). MIC reads that pass this filter should contain DNA sequences that are eliminated during rearrangement (IES, Internal Eliminated Sequence). The MIC reads covering MDS-IES breakpoints will split at MDS boundaries when mapped to MAC chromosomes (Figure 1B). SIGAR verifies that split read junctions indeed correspond to rearrangement breakpoints by searching within the split read for short sequence motifs, called “pointers”, which are microhomologous repeated sequences in the MIC that are retained as a single copy in the MAC after rearrangement (Figure 1A). SIGAR is able to infer more germline genome information from reads that map to two or more MDSs, even partially (Figure 1B). If the two blocks map adjacently to each other in the same direction, the region in between is recognized as a nonscrambled IES. Otherwise, the split read often indicates the presence of a scrambled region in the MIC.

### Validation of SIGAR by genome-assembly based annotations

To validate our proposed strategy to infer MIC genome features using short reads, we mapped *Oxytricha* Illumina MIC reads (Chen et al. 2014) to the original MAC genome assembly (Swart et al. 2013). SIGAR was then used to infer pointers from split reads (i.e. partially mapping reads), and the results were compared with existing MIC genome annotations (Chen et al. 2014; Burns et al. 2016). Pointer annotations in the MIC genome assembly were previously generated using MIDAS (Burns et al. 2016) which compares MIC and MAC genomes using BLAST. We find that 92.9% of the SIGAR-inferred pointers were validated by MIDAS. 59.70% contain the same first and last nucleotide in the MIDAS annotation, 27.74% share one identical boundary, and 3.8% share no boundary but have midpoints within 5bp of the MIDAS-inferred pointer (Table S1). The length distribution of SIGAR-inferred pointers is similar to that of MIDAS-inferred pointers, although fewer pointers were found in total (Figure 2A). In addition, SIGAR inferred a small number of “cryptic” pointers, defined as repeats longer than 20bp (Figure 2A) that differ from MIDAS-annotated MDS-MDS junctions. It is possible that some represent short regions of paralogy.

**Figure 2.**
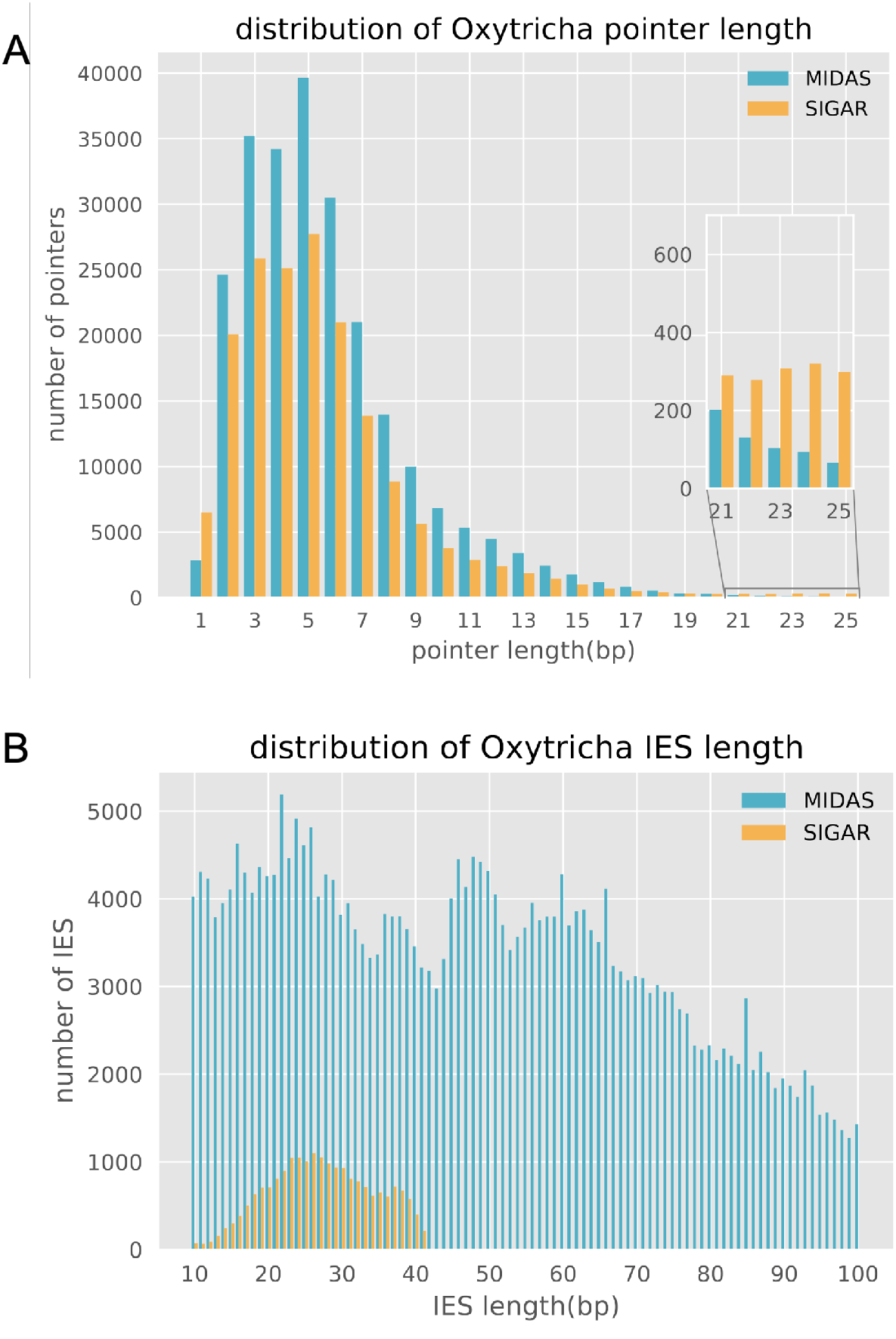
Comparison of pointer and IES length distributions between methods. **A)** The pointer length distribution and **B)** IES length distribution inferred for *Oxytricha* by MIDAS vs. SIGAR. Note that SIGAR used ~110X Illumina reads from Chen et al (2014), but MIDAS used an additional 15X PacBio reads for MIC genome assembly. Only ~60% of the MAC genome is considered by SIGAR as uniquely mapped regions for analysis, and the inferred IES length is restricted by read length (100bp in *Oxytricha* dataset). The pointer length distribution is only shown for 1-25bp, and IESs between 10-100bp.

SIGAR annotated half as many pointers as MIDAS (Table S1) because SIGAR was only applied to uniquely mapping regions of the MAC genome, in order to minimize the rate of false discovery. Such regions comprise 60.8-63.7% of the MAC genome, and contain 61.0-62.7% of all pointers (see Methods). SIGAR successfully annotated 80.4-82.3% of these pointers. We conclude that SIGAR recovers a large majority of pointers from the portion of the genome to which it was applied, even in the absence of a reference germline genome assembly.

In ciliates, the MIC DNA is significantly less abundant than MAC DNA, which can make it experimentally challenging to obtain pure micronuclei for DNA isolation. To test the robustness of SIGAR to variation in MIC coverage, we simulated 100 bp Illumina HiSeq reads from the *Oxytricha* MIC genome and calculated the number of inferred pointers. With only 5X MIC coverage, SIGAR was still able to recover 41.98% of MIDAS-inferred pointers, and 97.53% of SIGAR-inferred pointers were also identified by MIDAS (Table S1, Figure S1).

SIGAR can also infer the presence of short IESs that are contained within single Illumina reads (Figure 1B). Although the small read length constrains the size of IESs that can be detected from single reads, we were able to infer 20,599 short IESs with maximum length of 41bp using the 100bp read dataset (Figure 2B).

SIGAR’s results demonstrate that pointers can vary between alleles. 11,462 *Oxytricha* pointers validated by both MIDAS and SIGAR possess at least another pointer at the same junction inferred by SIGAR with high confidence (at least 2 reads supporting each boundary) (Table S2). Figure S2 highlights an example where pointer alleles correlate with SNPs in the MIC reads. Allele-specific information could only be detected by SIGAR (which examines individual sequencing reads) but not by assembly-based software like MIDAS, because genome assemblies tend to collapse alleles. This suggests that MAC chromosomes in *Oxytricha* arise from both MIC alleles, which can use different pointer sequences during MIC genome rearrangements.

### SIGAR infers scrambled genome architecture

Scrambled loci differ in order and/or orientation between the MIC and MAC versions (Figure 1A). Programmed rearrangements therefore entail MDS inversion and/or translocation. Scrambled genes have been described in most hypotrich ciliates (Chang et al. 2005; Hogan et al. 2001), *Chilodonella uncinata* (Katz and Kovner 2010) and some Postciliodesmatophora ciliates (Maurer-Alcalá et al. 2018). A recent report also validated the presence of scrambled loci in *Tetrahymena* (Sheng et al. 2020), providing further evidence that this is common to ciliate genomes. By using split read mapping, we find that SIGAR can infer some scrambled chromosome architectures and their associated pointers, even in the absence of a MIC genome assembly. When applied to *Oxytricha* datasets, SIGAR detected 8741 *Oxytricha* pointers at scrambled junctions. Figure 3 shows an example where SIGAR successfully inferred 3 out of 5 scrambled junctions in an *Oxytricha* MAC chromosome (Figure 3). SIGAR found 12 MIC reads covering both MDS1 and MDS3, while another 12 reads covered both MDS2 and MDS4 (Figure 3A and Figure S3).

**Figure 3.**
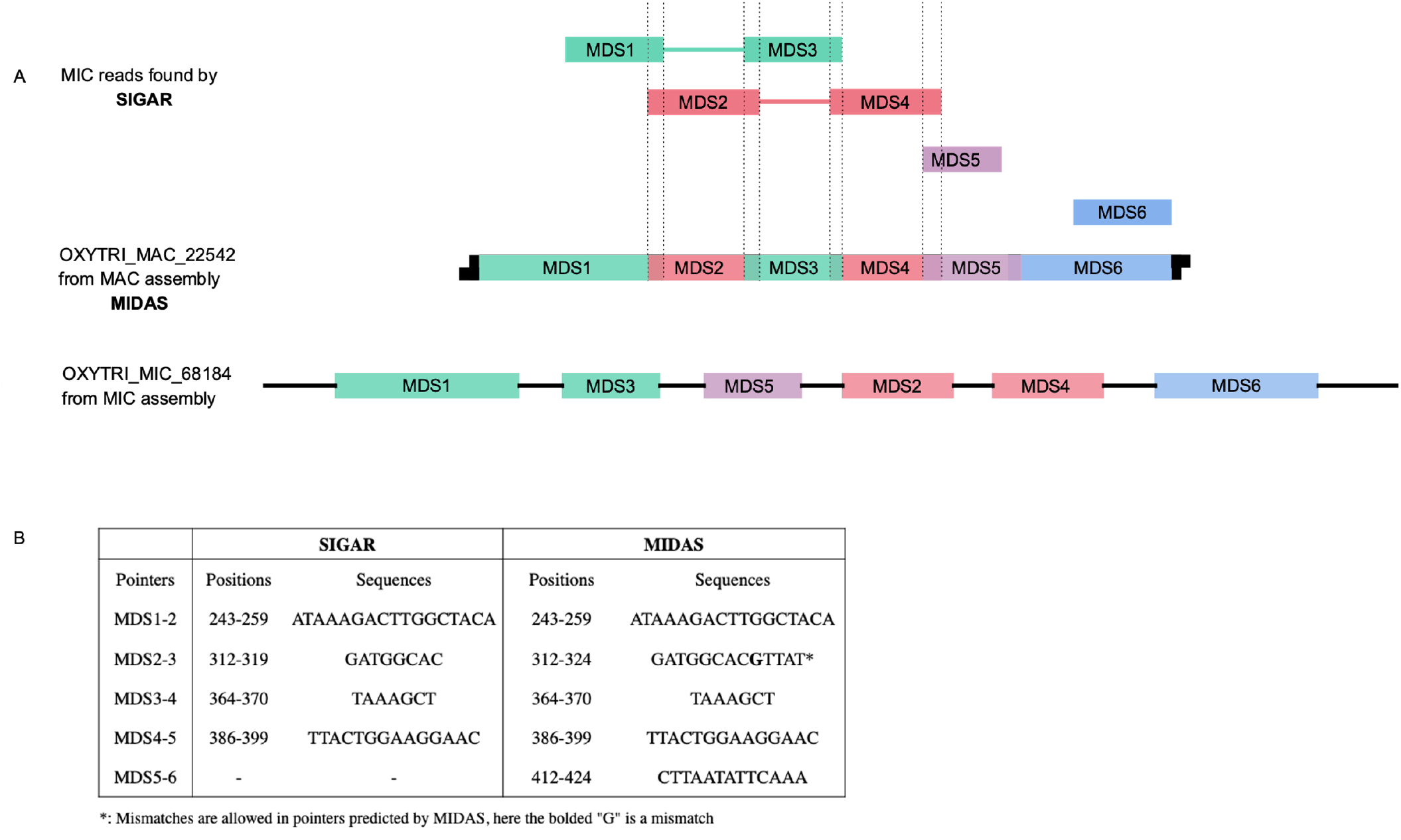
A representative scrambled region in *Oxytricha* inferred by SIGAR. **A)** SIGAR identified scrambled MIC reads that could be validated by MIDAS. The green, red, purple and blue reads are diagrams of split reads found by SIGAR mapped to OXYTRI_ MAC_22542 (see Figure S3 for reads mapping view). Green and red reads show a scrambled structure and the associated pointers are also inferred. **B)** The pointers on OXYTRI_ MAC_22542 found in SIGAR are the same as previous annotations by MIDAS.

### Inferring MIC genome architecture and rearrangement features in diverse ciliate species

SIGAR was developed to provide insights into germline rearrangement features across ciliates for which pure MIC DNA preparations are not readily available. We therefore applied SIGAR to 5 hypotrich genomes published in Chen et al. (2015), all of which have gene-sized MAC nanochromosomes like *Oxytricha*. Importantly, these sequencing datasets were derived from small numbers of whole cells, some from species that were not cultivated in the lab. We were able to infer pointers in all of these species using SIGAR (Table 1). TA is the most favored pointer among all 5 species. Though *Sterkiella* and *Urostyla* do not have TA as the most abundant pointer, both include A, T and TA as the three most abundant. With increasing evolutionary distance from *Oxytricha*, we observed more pointers containing TA as a substring. The GC content of short IESs (<0.2) is significantly lower than adjacent MDS regions (~0.3), consistent with surveys from other ciliate MIC genomes (Prescott 1994; Chen et al 2014). The pointer length distributions are similar in these species, except for *Sterkiella,* which exhibits an abundance of 5-20bp pointers (Figure S4).?

**Table 1.** Rearrangement features recovered by SIGAR for 5 surveyed hypotrich ciliates

| Phylogeny<br>(Chen et al.<br>2015) | Species | # of<br>pointers | Most<br>abundant<br>pointer | % of<br>"TA"<br>pointers | % of<br>pointers<br>with<br>"TA"<br>substring | # of<br>scrambled<br>pointers | #<br>of<br>IES | IES<br>G+C% | G+C%<br>of<br>MAC<br>contigs<br>with<br>IES | IES<br>G+C <<br>MDS<br>G+C<br>p-value<br>(t-test) |
| --- | --- | --- | --- | --- | --- | --- | --- | --- | --- | --- |
|  | *Oxytricha trifallax* |  |  |  |  |  |  |  |  |  |
|  | *Sterkiella histriomuscorum* | 12950 | A | 1.20 | 46.55 | 60 | 601 | 16 | 28 | 1e-69 |
|  | *Stylonychia lemnae* | 2509 | TA | 3.99 | 43.76 | 21 | 945 | 19 | 31 | 1e-120 |
|  | *Laurentiella sp.* | 1315 | TA | 5.10 | 48.14 | 19 | 509 | 14 | 28 | 1e-81 |
|  | *Paraurostyla sp.* | 1318 | TA | 7.28 | 52.66 | 17 | 653 | 11 | 30 | 1e-158 |
|  | *Urostyla sp.* | 1489 | A | 1.61 | 58.50 | 29 | 0 | - | - | - |

SIGAR also revealed evidence for novel scrambled loci in the 5 hypotrich ciliates (Table 1). Figure S5 shows the mapping view of an inferred scrambled locus in *Paraurostyla,* with pointers detected on each side. Short IESs were inferred for all species, except *Urostyla* (Table 1, Figure S4). Though short IESs were identified in *Urostyla* for *DNA pol α* (Chang et al. 2005) and *actin 1* (Hogan et al. 2001), we were unable to recover short IESs in the current dataset with limited MIC DNA. We note that intergenic IESs, which are common in *Tetrahymena* (Hamilton et al. 2016) cannot be detected by SIGAR, which identifies IESs adjacent to MDSs.

We also applied SIGAR to *Ichthyophthirius multifiliis (Ich),* an oligohymenophorean ciliate related to *Tetrahymena* (Coyne et al. 2011). *Ich* lives as a parasite in fish epithelia, causing “white spot” disease (Coyne et al. 2011). Furthermore, for ciliates that are hard to cultivate, SIGAR offers an ideal tool to infer properties of MIC genome architecture and DNA rearrangement, given the lack of availability of high quality MIC DNA preparations. Since the *Ich* MAC genome is ~84.1% A+T, we required pointers to have at least 5 well-mapped reads supporting each boundary. We found that the most abundant pointers are not only AT rich, but also contain surprisingly long TA tandem repeats (Figure S6). The pointer length distribution shows a peak at 10bp, with the 10bp pointer “TATATATATA” and “ATATATATAT” among the most abundant pointers in *Ich*.

## Discussion

Here, we have developed a novel computational tool, SIGAR, for inferring genome rearrangements and germline genome structure across diverse ciliates. A separate, complementary study proposed using short reads to infer the presence of eliminated DNA during genome rearrangement, but requires a high quality reference MIC genome assembly instead (Zheng et al. 2020). MIC genome assemblies typically pose the greatest challenge, whereas high quality MAC genome assemblies are much more accessible and also considerably less expensive to produce. In this sense, SIGAR will be more broadly applicable to the study of DNA rearrangements in ciliates.

SIGAR has provided new insights into the architecture of several ciliate MIC genomes, including *Ichthyophthirius multifiliis* and a group of early diverged hypotrichs, relative to *Oxytricha*. We observe a widespread preference for TA pointers across all ciliates in this survey. Studies of the ciliate model systems *Paramecium* and *Euplotes crassus* revealed the exclusive use of TA pointers in their germline genomes (Klobutcher and Herrick 1995; Arnaiz et al. 2012). It has also been shown that the terminal consensus sequences in *Paramecium* and *Euplotes* IES resemble terminal sequences in Tc1/mariner transposons (Klobutcher and Herrick 1995). This and other observations contributed to the hypothesis that an ancestral wave of transposons invaded ciliate germline genomes. The transposons then decayed but preserved the flanking “TA” as a relic and a modern requirement for accurate DNA elimination. Curiously, instead of this commonly observed 2bp TA pointer (Klobutcher and Herrick 1997, Chen et al. 2014), long TA repeats are present at *Ich* rearrangement junctions (Figure S6). Given that pointer sequences may help recruit enzymes that mediate IES removal and are necessary for excision in *Paramecium* (Aury et al. 2006), it is plausible that their extended length constitutes an adaptive feature to improve recognition and recruitment of DNA binding proteins that participate in genome rearrangement, amidst an AT-rich genome.

In addition to ciliates, many organisms in nature exhibit programmed genome rearrangement, such as lampreys (Smith et al. 2018) and songbirds (Kinsella et al. 2019). Furthermore, aberrant structural variations in mammalian cells are frequently observed in diseases like cancer (Stankiewicz and Lupski 2010; Forment et al. 2012). We expect SIGAR to be directly applicable to a wide range of genomes that exhibit rearrangements in both healthy and diseased states. SIGAR, which only requires one reference genome and short reads from a rearranged genome, could be a convenient tool to investigate all types of DNA rearrangement, providing insight into genome stability and instability.

## Methods

### SIGAR pipeline

SIGAR consists of three parts: 1) enrichment of MIC reads, 2) local alignment of MIC reads to MAC genome, and 3) parsing the split-read alignment report. At each step the pipeline provides adjustable parameters. All analyses in this paper were performed using the default parameters.

Step 1. We map input reads to MAC genome by Bowtie2 (Langmead and Salzberg 2012) end-to-end mapping to detect reads with only MDS. All reads with a mapping quality higher than threshold (default 3) are removed from the downstream analysis by SAMtools (Li et al. 2009).

Step 2. Filtered reads, which mainly consist of MIC reads are aligned to MAC contigs by BWA MEM local mapping (Li and Durbin 2009) with lowest mapping quality of 10 (default). MAC regions with abnormal high coverage, calculated by pileup.sh in BBtools (sourceforge.net/projects/bbmap/), were excluded from downstream analysis. The intermediate output of this step is used for visualization in this paper by IGV (Robinson et al. 2011).

Step 3. Parsing of the alignment output was implemented by Python. The main idea is parsing the CIGAR strings and “SA” tag in the alignment output. For example, CIGAR “40S60M” means that the initial 40 bp are soft clipped from mapping and a rearrangement junction is inferred at 40-41bp in the read. CIGAR “30M30I40M” means that a 30bp IES is inferred at a nonscrambled junction (Figure 1B). “SA” tags represent supplementary alignment of the read, indicating at least two mapping blocks present in a single read. These “SA” tagged reads are used to infer IES or scrambled loci.

Once SIGAR collects the split positions in reads by parsing CIGAR and “SA” tags, it infers pointers by pairwise comparison of alignments split in different directions (Figure 1B). Alignments are grouped as 5’ splits and 3’ splits. For example, “40S60M” is a 5’ split read, while “30M30I40M” contains a 30bp 3’ split alignment and a 40bp 5’ split alignment. The overlapped sequence between 5’ split and 3’ split is identified as a pointer. SIGAR outputs the pointers and the number of reads supporting each pointer boundary.

All source codes and manual for SIGAR are available at https://github.com/yifengevo/SIGAR.

### Genomes and datasets

The *Oxytricha trifallax* (strain JRB310) MAC genome data in this study are from MDS-IES-DB (http://knot.math.usf.edu/mds_ies_db/; Swart et al. 2013; Burns et al. 2016) and MIC Illumina reads are from GenBank SRX365496, SRX385993, SRX385994, SRX385995 and SRX385996 (Chen et al. 2014). Simulated MIC reads were produced by ART (Huang et al. 2012). To estimate uniquely mapped regions in SIGAR analysis, we simulated 30bp and 92bp reads using MAC genome as reference, to mimic partially aligned regions in split reads, representing the minimum and maximum SIGAR split reads alignment. We mapped these reads to MAC genome by BWA MEM and filtered by mapq 10, the default setting of SIGAR. 63.69% reference bases are covered for 92bp reads and 60.77% for 30bp reads, which indicates that ~60% MAC genome is considered in the SIGAR analysis.

The hypotrich MAC genomes and whole cell DNA sequences are from Chen et al. 2015, accession numbers *Laurentilla sp.* LASS02000000, *Sterkiella histriomuscorum* LAST02000000, *Stylonychia lemnae* ADNZ03000000, *Urostyla sp.* LASQ02000000, and *Paraurostyla sp.* LASR02000000. Only telomeric MAC contigs with “CCCCAAAACCCC” or “GGGGTTTTGGGG” were used in the analysis. All SIGAR annotations are within the contig body, which is at least 50bp from contig ends to avoid noisy mapping on telomeric regions.

*The Ichthyophthirius multifiliis* MAC genome is from Coyne et al. 2011, Genbank accession number GCF_000220395.1. *Ichthyophthirius multifiliis* whole cell DNA sequence reads were from MacColl et al. 2015.

## Acknowledgements

We thank Theodore Clark and Donna Cassidy-Hanley for sharing *Ich* data and Sindhuja Devanapally for comments on the manuscript. We thank Rafik Neme, Jananan Pathmanathan, Talya Yerlici, Derek Clay, Sandrine Moreira, and all laboratory members for discussion. We are grateful for the National Center for Genome Analysis Support (NCGAS) computing resources (supported by NSF DBI-1062432, ABI-1458641, ABI-1759906 to Indiana University). This work was supported by NIH grant R35GM122555 and NSF grant DMS1764366 to L.F.L.

