## Supplementary materials for "SIGAR: Inferring features of genome architecture and DNA rearrangements by split read mapping"

**Table S1.** SIGAR-inferred pointers compared to MIDAS annotation

| Pointers | Real Illumina reads  (~110X, Chen et al. 2014) | Simulated Hiseq2000 dataset 5X 100bp | Simulated Hiseq2000 dataset 20X 100bp | Simulated Hiseq2000 dataset 20X 100bp (split reads freq>=2, high confidence set) |
| --- | --- | --- | --- | --- |
| Total number | 184662 | 135849 | 249360 | 134277 |
| Same start and end | 59.70% | 67.07% | 43.11% | 79.05% |
| One identical boundary | 27.74% | 26.85% | 42.93% | 16.88% |
| Similar but not identical boundary (mid point ±5bp) | 3.80% | 3.12% | 10.12% | 1.54% |
| Vaguely inferred by MIDAS but resolved by SIGAR | 1.67% | 0.50% | 0.41% | 0.47% |
| SIGAR specific^a^ | 7.10% | 2.47% | 3.84% | 2.53% |
| Ratio of SIGAR-inferred pointers in all MIDAS pointers | 50.47% | 41.98% | 48.74% | 48.22% |
| Ratio of SIGAR-inferred pointers in MIDAS pointers within SIGAR-applied^b^ regions | 80.4-82.3% | 67.0%-68.8% | 77.7-79.8% | 76.8-79.0% |

^a^ Pointers not found in MIDAS

^b^ See methods for the uniquely mapped regions used in SIGAR.

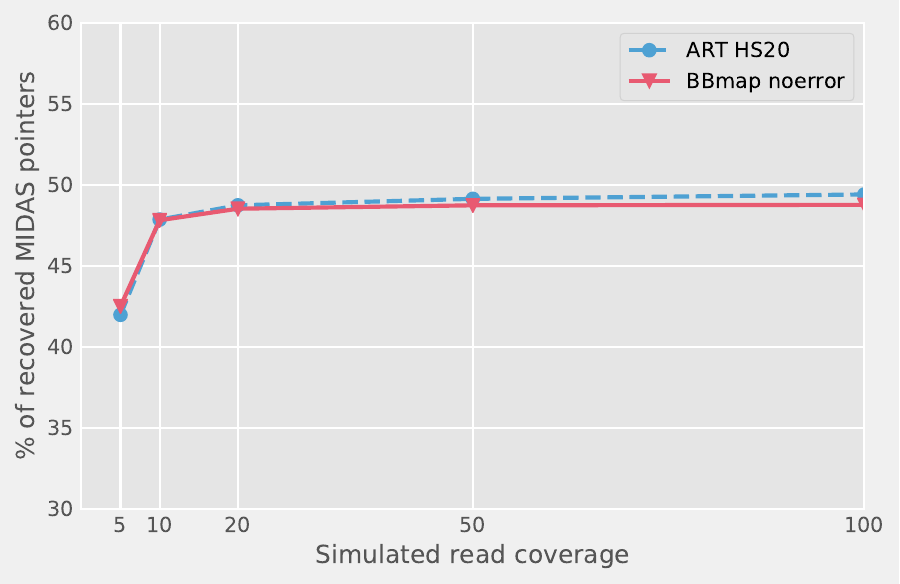

**Figure S1.** The ratio of recovered MIDAS pointers using simulated *Oxytricha* MIC reads as input of SIGAR. We used two tools (ART and BBmap) to simulate 5X, 10X, 20X, 50X and 100X 100bp reads from the *Oxytricha* MIC genome assembly. ART simulated Illumina Hiseq2000 reads with “-ss HS20”. BBmap randomreads.sh was used to simulate error-free reads.

**Table S2.** Allele specific pointers identified by SIGAR in *Oxytricha trifallax* (strain JRB310)

| Contig | start | end | # of Reads mapping to MDS n | # of Reads mapping to MDS n+1 | pointer |
| --- | --- | --- | --- | --- | --- |
| OXYTRI_MAC_5 | 1377 | 1383 | 36 | 61 | TAAACAT |
| OXYTRI_MAC_5 | 1379 | 1383 | 9 | 61 | AACAT |
| OXYTRI_MAC_7 | 5080 | 5086 | 93 | 67 | ATATTTT |
| OXYTRI_MAC_7 | 5080 | 5087 | 93 | 15 | ATATTTTT |
| OXYTRI_MAC_7 | 5430 | 5435 | 34 | 29 | AACCAA |
| OXYTRI_MAC_7 | 5430 | 5436 | 34 | 15 | AACCAAT |
| OXYTRI_MAC_9 | 217 | 220 | 46 | 52 | AGAA |
| OXYTRI_MAC_9 | 217 | 221 | 46 | 15 | AGAAT |
| OXYTRI_MAC_9 | 218 | 220 | 7 | 52 | GAA |
| OXYTRI_MAC_9 | 218 | 221 | 7 | 15 | GAAT |
| OXYTRI_MAC_18 | 2019 | 2023 | 80 | 24 | TAATA |
| OXYTRI_MAC_18 | 2019 | 2027 | 80 | 32 | TAATATATC |

Shaded rows are the common pointers annotated by both MIDAS and SIGAR.

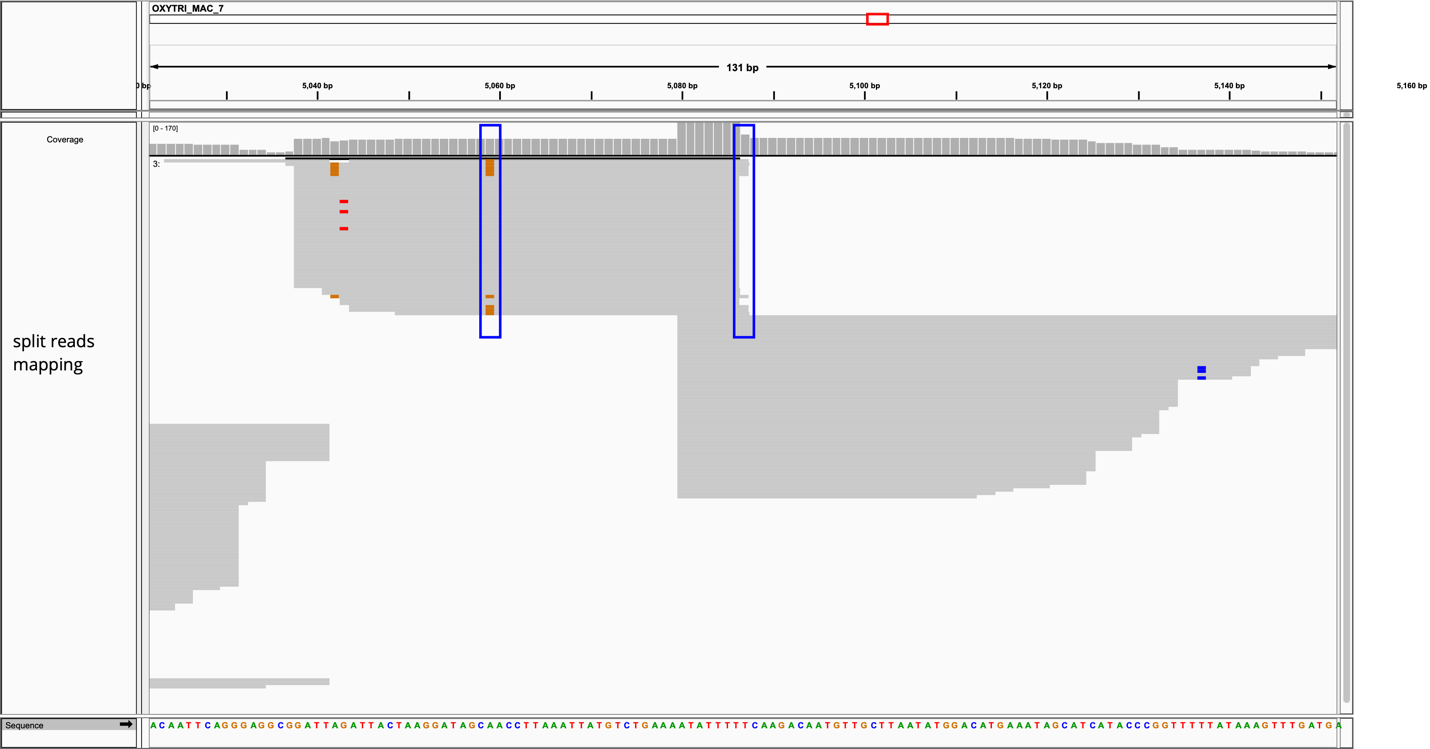

**Figure S2.** Identified pointer alleles correlate with SNPs in MIC reads. The pointer sequence is **ATATTTT** (position 5080-5086) when the split reads have an “A” at position 5059. On the other hand, the pointer sequence is ATATTTT**T** (position 5080-5087) when the split reads have a “G” at position 5059 (highlighted in blue box with brown color). The number of split reads of each allele is shown in Table S2 for OXYTRI_MAC_7.

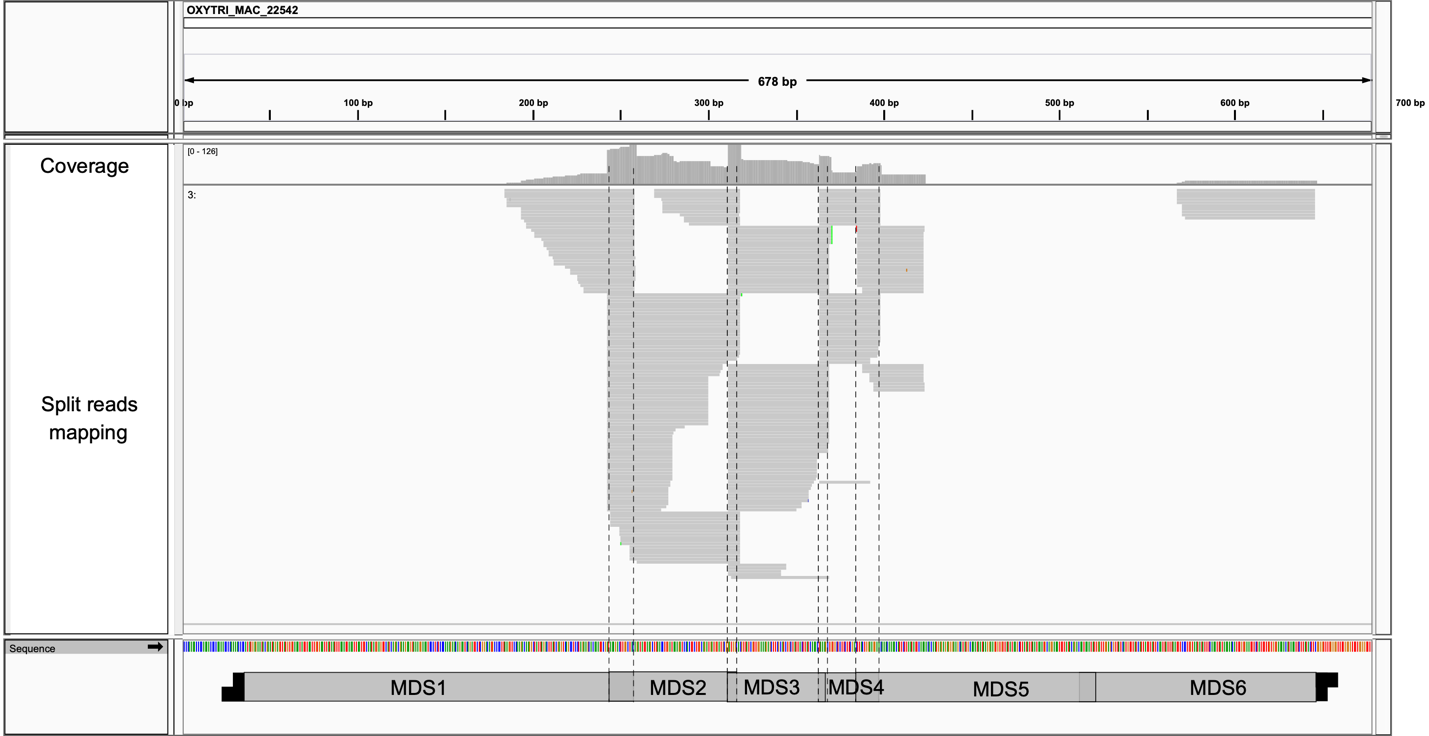

**Figure S3.** The mapping view of a scrambled *Oxytricha* MAC chromosome. The diagram of this mapping view is shown in Figure 3. The dashed lines highlight boundaries of MDS inferred by SIGAR.

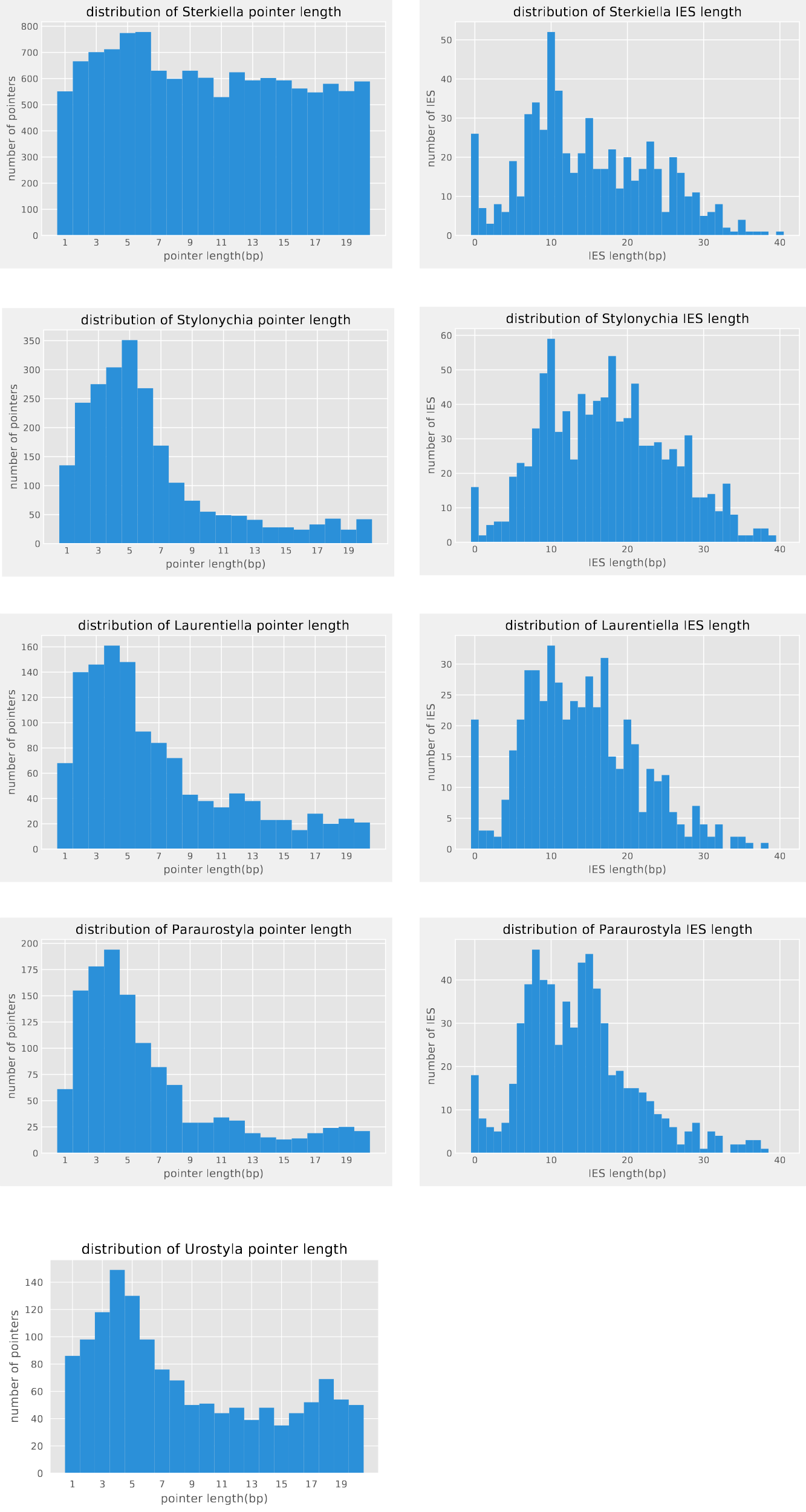

**Figure S4.** The pointer and IES length distribution of 5 hypotrich ciliates inferred by SIGAR. No short IESs were identified in the *Urostyla* data.

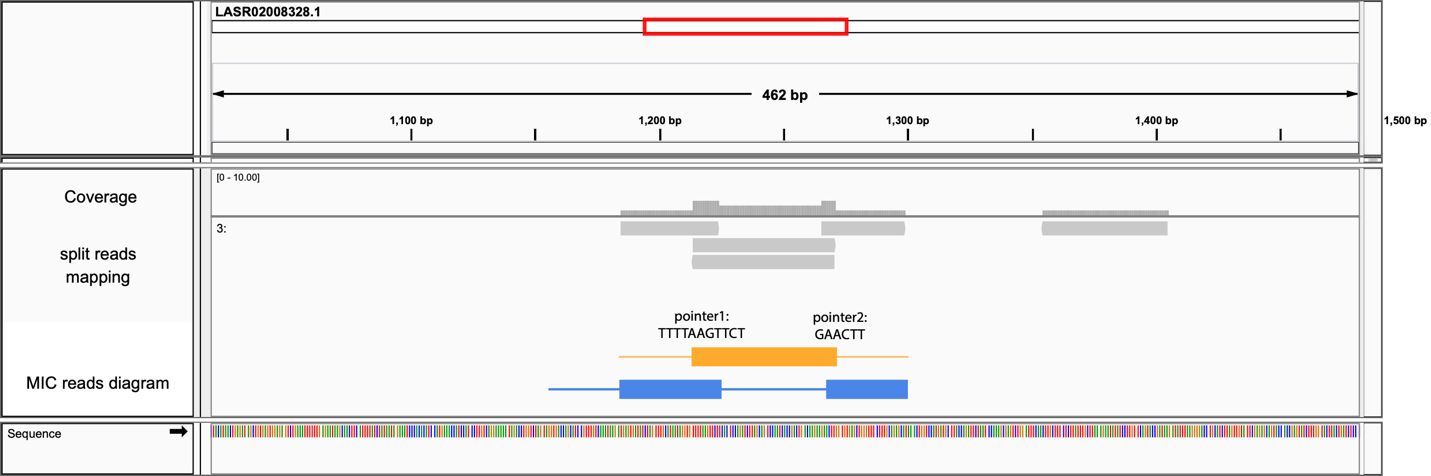

**Figure S5.** A scrambled example in *Paraurostyla* *sp*. inferred by SIGAR. The yellow and blue reads are diagrams of mapped split reads. The pointer sequences at two junctions are also inferred.

| Most abundant Ich pointers | |
| --- | --- |
| pointer | frequency |
| A | 118 |
| T | 112 |
| TA | 89 |
| ATA | 75 |
| TAT | 70 |
| ATATATATAT | 63 |
| AT | 55 |
| ATAT | 52 |
| ATATA | 40 |
| ATATATAT | 38 |
| ATATATA | 37 |
| TATAT | 33 |
| ATATAT | 31 |
| ATATATATATAT | 31 |
| TATATATATA | 27 |
| TATATATAT | 27 |

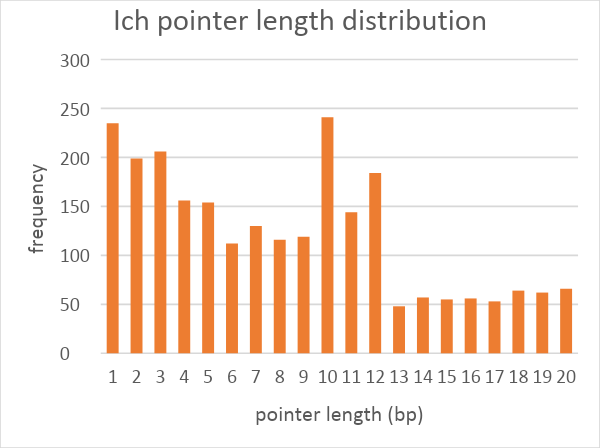

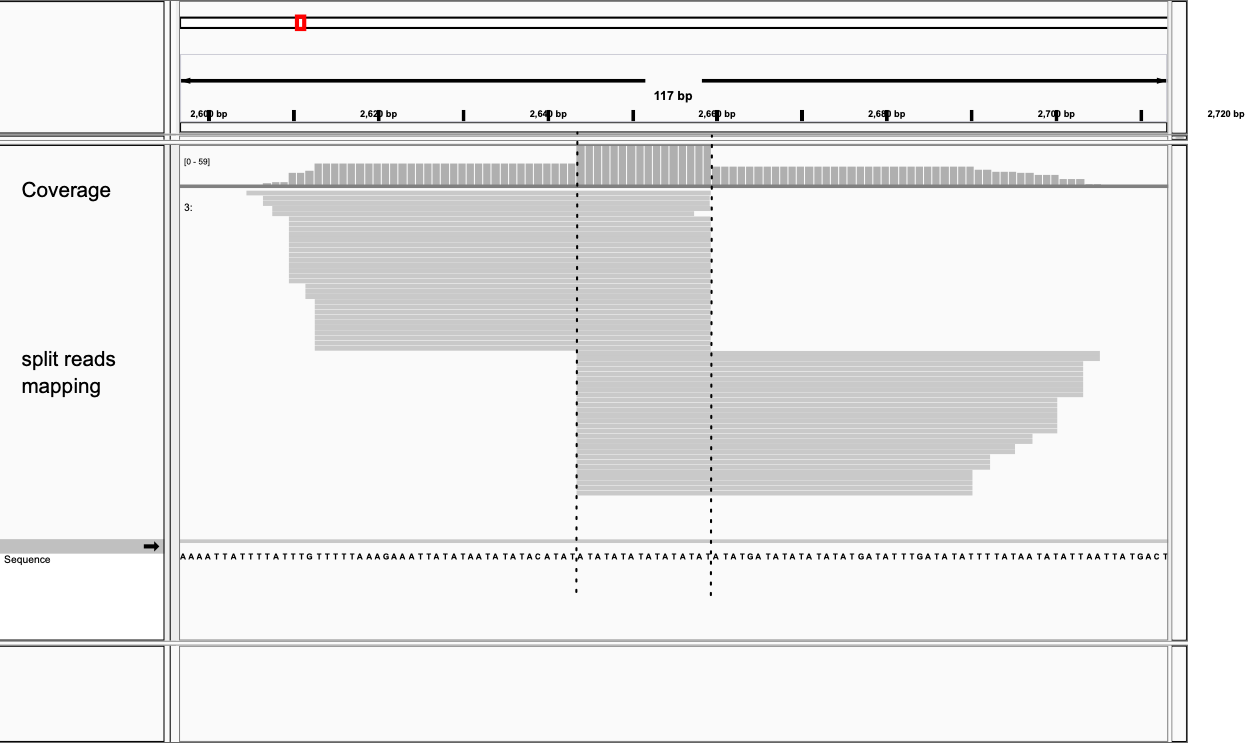

**Figure S6.** *Ichthyophthirius multifiliis (*Ich*)* pointers inferred by SIGAR. A) Most abundant pointers. B) Pointer length distribution. C) A representative mapping view of a SIGAR-inferred pointer. The pointer is at 2643-2659bp on MAC contig NW_004086212.1, supported by 28 and 30 split reads on each adjacent MDS.
